# Propagating activity in neocortex, mediated by gap-junctions and modulated by extracellular potassium

**DOI:** 10.1101/669226

**Authors:** Christoforos A. Papasavvas, R. Ryley Parrish, Andrew J. Trevelyan

## Abstract

Parvalbumin-expressing interneurons in cortical networks are coupled by gap-junctions, forming a syncytium that supports propagating epileptiform discharges, induced by 4-aminopyridine. It remains unclear, however, whether these propagating events occur under more natural states, without pharmacological blockade. In particular, we investigated whether propagation also happens when extracellular K^+^ rises, as is known to occur following intense network activity, such as during seizures. We examined how increasing [K^+^]_o_ affects the likelihood of propagating activity away from a site of focal (200-400µm) optogenetic activation of PV-interneurons. Activity was recorded using a linear 16-electrode array placed along layer V of primary visual cortex. At baseline levels of [K^+^]_o_ (3.5mM), induced activity was recorded only within the illuminated area. However, when [K^+^]_o_ was increased above a threshold level (50^th^ percentile= 8.0mM; interquartile range= 7.5-9.5mM), time-locked, fast-spiking unit activity, indicative of parvalbumin-expressing interneuron firing, was also recorded outside the illuminated area, propagating at 59.1 mm/s. Blockade of glutamatergic synaptic transmission reduced the efficacy of propagation, but could be restored by further increasing [K^+^]_o_. Propagation was further reduced, and in most cases prevented altogether, by pharmacological blockade of gap-junctions, achieved by any of three different drugs, quinine, mefloquine or carbenoxolone. Wash-out of quinine rapidly re-established the pattern of propagating activity. Computer simulations show qualitative differences between propagating discharges in high [K^+^]_o_ and 4-aminopyridine, arising from differences in the electrotonic effects of these two manipulations. These interneuronal syncytial interactions are likely to affect the complex electrographic dynamics of seizures, once [K^+^]_o_ is raised above this threshold level.

**Significance statement:** We demonstrate the spatially extended propagation of activity through a gap-junction mediated syncytium of parvalbumin-expressing interneurons, in conditions that are known to exist at times within the brain. Previous work has only shown gap-junction coordination very locally, through directly connected cells, or induced at a distance by pharmacological means. We show that cell-class specific spread is facilitated by raised extracellular K^+^. This is highly pertinent to what happens at the onset of, and during, seizures, when extracellular K^+^ can rise rapidly to levels well in excess of the measured threshold for propagation. Our data suggests that interneuronal coupling will be enhanced at this time, and this has clear implications for the behaviour of these cells as seizures progress.

## Introduction

Cortical interneurons are connected by gap-junctions to other interneurons within the same class (Galarreta & Hestrin, 1999; Gibson *et al*., 1999; Galarreta & Hestrin, 2001; Amitai *et al*., 2002; Hestrin & Galarreta, 2005; Juszczak & Swiergiel, 2009), providing a highly specific, excitatory link between these cells. Initial studies showed that gap-junction coupling normalised the voltage difference between two cells and further synchronised their firing with millisecond precision (Galarreta & Hestrin, 1999; Gibson *et al*., 1999; Bennett & Zukin, 2004; Connors & Long, 2004). These studies, however, only examined very localised sets of interneurons, and the question of how widely this synchronisation spreads remained unexamined.

Propagation through this gap-junction coupled network has been observed in one particular pathological condition, in a widely used model of epileptic activity, induced by bathing brain slices in 4-aminopyridine (4-AP) (Szente *et al*., 2002; Gajda *et al*., 2003; Gigout *et al*., 2006a). 4-AP blocks various voltage-dependent K^+^ channels, and appears to have a disproportionate effect on the population of fast-spiking interneurons, inducing rhythmic bursting in these cells even in the absence of any glutamatergic drive (Avoli *et al*., 2002; Bohannon & Hablitz, 2018; Parrish *et al*., 2018). These epileptiform discharges propagate reliably across brain slices, with a broad and relatively slow wave-front. This though represents a rather specific case, in which the spread of activity through the syncytium is facilitated by two factors related to the K^+^ conductance blockade, with cells being depolarised and also electrotonically more compact, and which may not occur *in vivo*. In contrast, we speculated that gap-junction mediated propagation may also be supported by a different change in the neuronal milieu, that has been demonstrated both *in vivo* and *in vitro*: namely, the raised extracellular potassium ([K^+^]_o_) associated with extreme levels of neuronal activity. We now show that this is indeed the case, although the pattern of spreading activity differs qualitatively from that seen in 4-AP. We used mice that express channelrhodopsin under the PV promoter, allowing us to stimulate specifically this population of interneurons in a highly focal manner. We then examined the time-locked activity propagating out from this focus of activation. We show that this propagation is sensitive to, but not dependent on, glutamatergic synaptic activity, but it is abolished by any of three different gap-junction blockers. We then explore the differences between the two patterns of spreading activity, induced either by high [K^+^]_o_ or 4-AP, using an extended, multineuron, compartmental model of the interneuronal syncytium.

## Methods

### Cortical expression of optogenetic proteins

All animal handling and experimentation were done according to the guidelines laid by the UK Home Office and Animals (Scientific Procedures) Act 1986 and approved by the Newcastle University Animal Welfare and Ethical Review Body (AWERB # 545). Cortical channelrhodopsin-2 (ChR2) expression was achieved by using genetically engineered transgenic mice. Brain slices were prepared from first generation cross-breeding of homozygous floxed-channelrhodopsin mice (B6; 129S-Gt(ROSA)26Sortm32(CAG-COP4*H134R/EYFP)Hze/J; Jackson Laboratory, stock number 012569) with homozygous PV-cre mice (B6; 129P2-Pvalbtm1(cre)Arbr/J; Jackson Laboratory, stock number 008069).

### Preparation of brain slices

Twenty-two young mice (6-12 weeks old) were sacrificed to prepare the brain slices. They were first anesthetised using Ketamine (0.3 mL / 30 g) and then perfused with ice-cold sucrose-based artificial cerebrospinal fluid (ACSF: NaHCO3 24mM, KCl 3 mM, NaH2PO4 1.25 mM, sucrose 227.8 mM, glucose 10mM, MgCl2 4 mM) before the brain was removed to prepare coronal brain slices (400μm thick). The slices were cut on Leica VT1200 vibratome (Leica Microsystems, Wetzlar, Germany) in ice-cold oxygenated (95% O2/ 5% CO2) sucrose-based ACSF (NaCl 125mM, NaHCO3 26mM, glucose 10mM, KCl 3.5 mM, NaH2PO4 1.26 mM, MgCl2 3mM). After cutting, the slices were transferred to an incubation, interface chamber (room temperature) perfused with oxygenated (normal) ACSF (NaCl 125 mM, NaHCO3 26 mM, glucose 10 mM, KCl 3.5 mM, NaH2PO4 1.26 mM, CaCl2 2 mM, MgCl2 1 mM) for at least 1 hour before transferring them to a recording interface chamber. In the recording interface chamber, the ACSF used was normal ACSF as above. The ACSF was perfused at 1.5 2.5 ml /min and its temperature was kept at 33-36°C. In the case of the ChR2 experiments, the concentration of extracellular K+ was systematically increased during the recording by adding KCl to the perfused ACSF.

### Extracellular recordings

Multichannel extracellular recordings were collected at 25 kHz unless otherwise stated, using a linear 16-channel-probe configuration (A16×1-2mm-100-177; NeuroNexus; electrode separation, 100μm). This was connected to an ME16-FAI-μPA system and MC-Rack software (Multichannel Systems, Reutlingen). The signals were filtered using an analog high-pass filter with a 300Hz cutoff frequency. Data acquisition was carried out using a 1401-3 Analog-Digital converter (Cambridge Electronic Design, Cambridge) and Spike2 software (Cambridge Electronic Design, Cambridge). The electrode array was placed along layer V in the occipital dorsal area of neocortex, approximately corresponding to primary visual cortex (Dong, 2008). Channelrhodopsin was activated by a 470nm LED, delivering light through a Nikon Plan Fluor 4x objective (NA 0.13), using the patterned illuminator Polygon400 (Mightex Systems, Pleasanton, CA, USA). The system was controlled and the patterns were designed through the PolyScan 2 software from the same company. The light intensity was measured at approximately 2mW/mm^2^.

AMPA currents were blocked by bathing in 20μM NBQX (HelloBio) and NMDA currents were blocked using 50μM d-APV (Abcam Biochemicals). We used three different gap junction blockers, namely Mefloquine (50μM), Quinine (100μM, both from Sigma-Aldrich, Gillingham, UK) and Carbenoxolone (100μM, Tocris Bioscience, Bristol, UK).

### Simulations

The model cell consists of three compartments: one soma (27μm length, 29μm diameter, 1 segment) and two morphologically and biophysically identical dendrites (left and right; 200μm length (Fukuda *et al*., 2006), 0.8μm diameter (Fukuda & Kosaka, 2003), 10 segments). The interneuron model used in this study is adapted from the biophysically detailed interneuron model from Konstantoudaki *et al*., 2014 (ModelDB, Accession number: 168310). The soma is equipped with the following mechanisms: fast Na+ current, A-type K+ current, delayed-rectifier K+ current, slow K+ current, N-type high-threshold activated Ca++ current, hyperpolarization-activated cation current (Ih), fast afterhyperpolarization K+ current and a Ca++ buffering mechanism. The dendrites are equipped with the following mechanisms: fast Na+ current, A-type K+ current, and delayed-rectifier K+ current. The conductance values used are shown in Table 1. Seventy identical cells are scattered in a virtual slice with dimensions 650 x 150 x 150μm. Thus, the density of the population is approximately 5000 PV interneurons / mm^3^, which is close to the density found in layer V of mouse primary visual cortex (Pakan *et al*., 2016). The cells were randomly connected with gap junctions following the connectivity rule in Fig. 8B. A pair of cells would have a maximum of one gap junction (g_gap_ = 0.3nS; Fukuda *et al*., 2006; Galarreta & Hestrin, 2002) connecting the right dendrite of the cell on the left with the left dendrite of the cell on the right. The placement of the gap junction along the dendrites was symmetric and was randomly placed with a uniform probability distribution between the closest point possible (considering the distance between the cells) and the full length of the dendrite. For example: two cells with 100μm distance between them could be connected from the 50μm dendritic point away from their soma up to the most distal dendritic point, that is, 200μm away from their soma.

**Table 1:** Biophysical mechanisms used in the model and their conductance values.

| mechanism | soma | dendrite |
| --- | --- | --- |
| Na<sup>+</sup> conductance (S/cm<sup>2</sup>) | 0.045 | 0.06 |
| Delayed rectifier K<sup>+</sup> | 0.018 | 0.009 |
| N-type Ca<sup>++</sup> | 0.0003 | N/A |
| D-type K<sup>+</sup> | 0.0000725 | N/A |
| H-current | 0.00001 | N/A |
| A-type K<sup>+</sup> | 0.048 | 0.48 |
| fAHP | 0.0001 | N/A |
| Ca<sup>++</sup> diffusion | yes | no |
| Cm (μF/cm<sup>2</sup>) | 1.2 | 1.2 |
| Ra (ohm/cm) | 150 | 150 |
| Rm (kOhm cm<sup>2</sup>) | 10 | 10 |

The cells that were located in the leftmost 200μm of the virtual slice were directly stimulated at the soma with a 25ms pulse of 0.55nA amplitude starting at 50ms into the simulation. The speed of propagation is calculated based on the first propagation wave, that is, the first spikes of two specific cells. It is equal to the distance travelled over time between the last cell in the stimulation area and the last cell in the overall propagation. For the results shown in Fig. 8E and 8F, only the first 200 long propagations were considered for the analysis. A long propagation is considered a propagation that reaches the 60^th^ cell or beyond that.

All simulations were run using the NEURON simulator (Hines & Carnevale, 2001) through Python (PyNN interface (Davison *et al*., 2008)).

## Results

### Optogenetic activation of parvalbumin-expressing cells

We investigated the propagation of activity through the syncytium of PV-expressing interneurons, in occipital cortical brain slices in different levels of extracellular K^+^. Extracellular recordings were made from 53 mouse brain slices prepared from 22 young adult mice which expressed ChR2 under the PV promoter. We recorded extracellular field potentials using a linear multi-electrode array (MEA; 1.5 mm wide array of 16 electrodes with 0.1 mm spacing between the shafts) placed along layer V (Fig. 1B), where there is a dense network of electrically coupled PV-expressing interneurons (Galarreta & Hestrin, 1999; Gibson *et al*., 1999; Fukuda & Kosaka, 2003). We optogenetically activated these PV-expressing interneurons specifically, in a small and circumscribed area, extending over 3-4 adjacent electrodes, using a focused patterned illuminator (typically 300-400 x 100 μm, see Fig. 1C and Methods). The blue light was delivered as a train of pulses lasting 3s, at 20 Hz frequency with 50% duty cycle, and repeated every 20s. The spread of activity beyond the light spot was assayed using a linear multielectrode array, which typically extended at least 0.7 mm beyond the light spot, sampling at 100µm spaces between individual electrodes.

**Figure 1.**
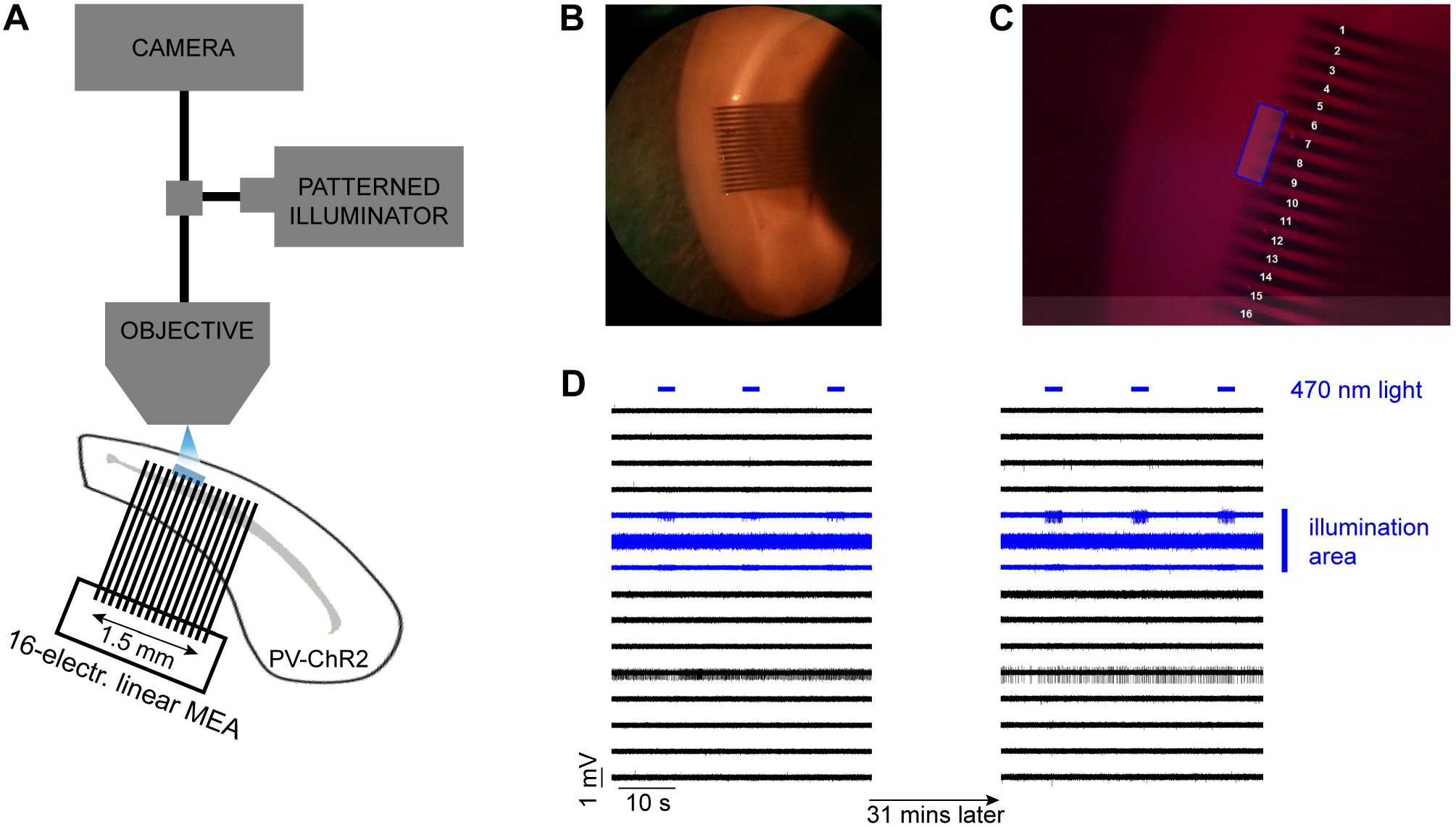
Schematic of the experimental setup and an example of the control experiment. (A) Extracellular recordings were taken using a 16-electrode linear multi-electrode array (MEA) placed along layer V in a mouse cortical brain slice in which ChR2 was expressed in PV^+^ cells. The distance between adjacent electrodes was 100 μm. Photostimulation (blue LED) was delivered using a patterned illuminator through the microscope objective. (B) The recordings were taken from the dorsal area of the slice, targeting the primary visual area. (C) Example of an illumination pattern (4-electrode wide area) which was drawn in the middle of the array using the patterned illuminator software. (D) A typical control experiment during which repeated illumination was delivered for 31 minutes while the extracellular K^+^ remained at normal levels. The evoked activity was only seen at electrodes within the illumination area, and there was no propagating activity, a pattern that remained stable for the entire duration of the recording (31 mins).

### Propagation of activity with increased extracellular K^+^

At baseline levels of extracellular K^+^ ([K^+^]_o_ = 3.5mM), neuronal spiking was reliably recorded at those electrodes only within the spot of light, or occasionally from electrodes immediately adjacent, presumably from instances where cells had processes extending into the illuminated area. Critically, though, in this baseline condition, we never recorded triggered activity beyond this restricted site. This pattern of spatially restricted, time-locked activity was very stable, when [K^+^]_o_ was kept constant for an extended period (>30mins, n = 3 brain slices, 90 stimulations; Figure 1D). We then increased [K^+^]_o_, with increments of 1-2mM every 5-10 mins, to investigate how this affected activity patterns. Predictably, this produced a marked increase in spontaneous activity (Korn *et al*., 1987; Jensen & Yaari, 1997), and additionally, in 45.3% of all brain slices (24 out of 53 brain slices) we also noted bursts of firing in electrodes away from the illumination site (at least 150μm away), that were time-locked, at short latencies, to the pattern of illumination. Notably, this transition happened rapidly, typically within 2mins, after a threshold level of [K^+^]_o_ was reached, and then remained stable thereafter. Thus, the raised extracellular [K^+^] facilitated the spread of activity away from the focal site of optogenetic activation (Fig. 2).

**Figure 2.**
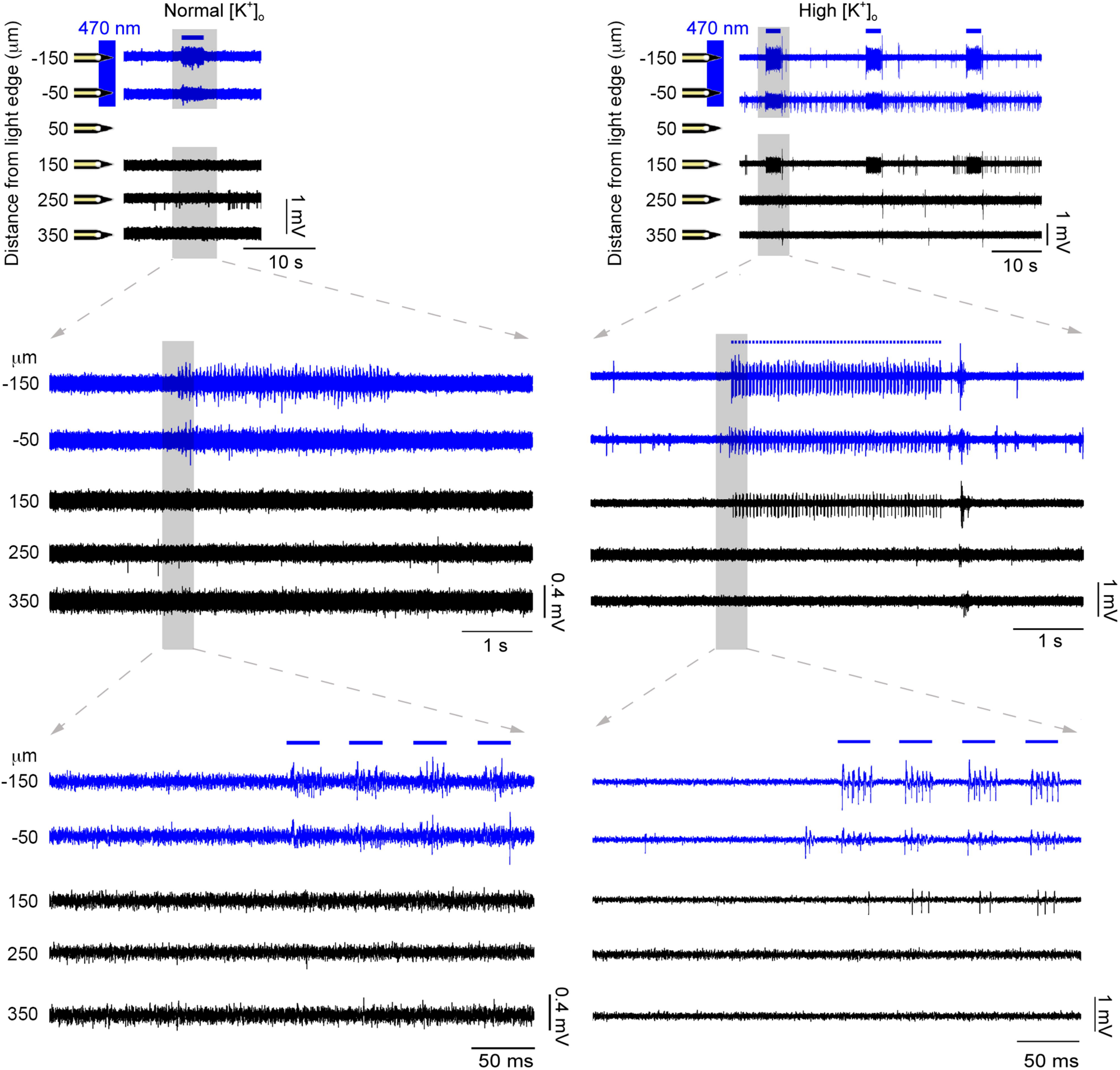
Propagating activity arising with raised extracellular K^+^. The area around a subset of the electrodes was illuminated (marked with blue) with 3-second long pulse trains (20Hz, 50% duty cycle), repeated every 20 s. PV^+^ cells around these electrodes responded with 4-5 spikes per pulse (see zoomed-in traces). After raising the extracellular K^+^ concentration (from 3.5mM to 7.5mM), induced activity was recorded in a distant electrode as well (150μm away). The activity recorded outside the illumination area was time-locked to the activity inside that area, albeit with a short delay. Considering that the photostimulated cells are GABAergic in nature, the activity at the distant electrode is hypothesized to propagate through the electrical synapses between PV^+^ cells.

The proportion of brain slices that showed propagation away from the point of illumination increased, as [K^+^]_o_ was raised (Fig. 3). Considering only those slices which showed propagation, we fitted a sigmoidal curve to this data, to derive the threshold level of [K^+^]_o_ supporting propagation (50% of trials). The threshold was 8.0 +/− 0.15 mM (median +/− 95% confidence interval; Interquartile range = 7.5-9.5mM; full range = 4.5-11.5mM).

**Figure 3.**
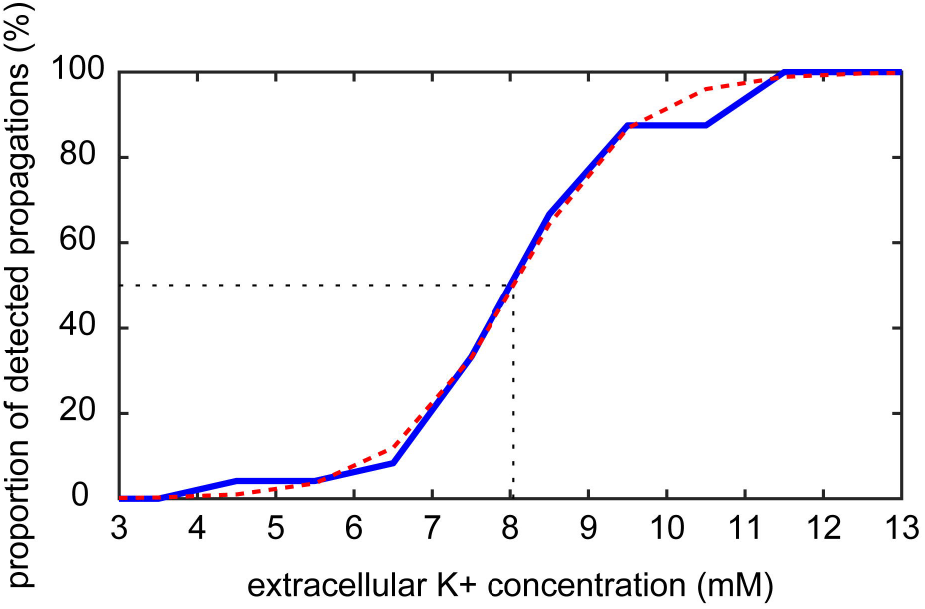
Propagation is facilitated by raised extracellular K^+^. The plot shows the cumulative proportion of detected propagations at different levels of extracellular K^+^ concentration. The majority of propagations (16 out of 24, 66.7%) were observed with an increase of the extracellular K^+^ up to 8.5 <u>mM</u>. The threshold (at the 50% point) was calculated at 8.0 mM after fitting a sigmoidal function (red dashed curve).

### Propagating activity was sensitive to glutamatergic and gap-junction blockers

We hypothesised that the propagating activity spread through gap-junction connections between PV-expressing interneurons. Examining only brain slices that showed clear propagation of activity beyond the illumination site (20 slices), and working at increased [K^+^]_o_ (mean = 8.40mM; range = 4.5-11.5mM), we first blocked glutamatergic currents using antagonists of AMPA and NMDA receptors (20µM NBQX and 50µM D-APV, respectively), to assess what contribution, if any, was made through conventional synaptic excitatory pathways. This also served to reduce the level of spontaneous activity. In a proportion of brain slices (6 out of 20 slices), following the introduction of glutamate blockers, the evoked propagating activity gradually diminished in parallel with the reduction in spontaneous activity, indicating that synaptic excitation did indeed contribute to these events (Figures 4 and 5). Notably, we were able to resurrect the propagating event by further increasing the extracellular K^+^ (Figure 5), indicating that propagation is facilitated by glutamatergic activity within the slice, but is not dependent on it. In the remaining brain slices (14/20 slices), propagation of activity persisted, following glutamatergic blockade, and indeed was more apparent because it existed on a lower level of background activity.

**Figure 4.**
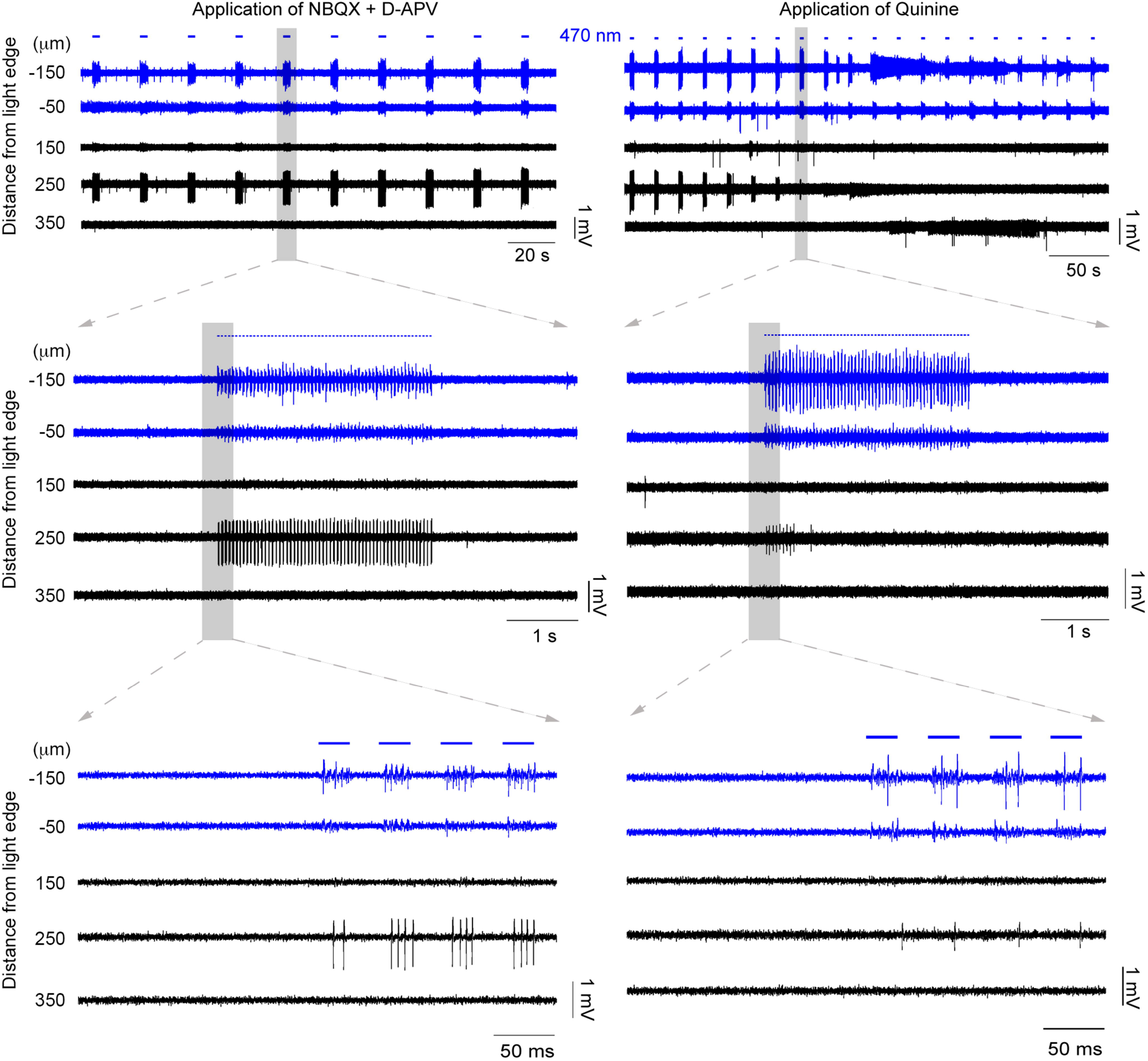
Propagating activity is sensitive to drugs that block glutamatergic receptors and also gap-junction blockers. The underlying mechanisms of the propagating activity were investigated by the application of pharmacological agents. This example is the continuation of the example in Fig. 2. First, the glutamate receptors were blocked by applying NBQX and D-APV (left panel). In this example, the spontaneous activity was decreased but the propagated activity at the distant electrode remained strong. This indicates that glutamate release (e.g., through disinhibition) was not involved in the propagation. Then, the gap junction blocker quinine was applied (right panel). The activity, recorded 250μm away, was gradually suppressed, and eventually silenced altogether, as evident from the decreasing amplitude and spike rate.

**Figure 5.**
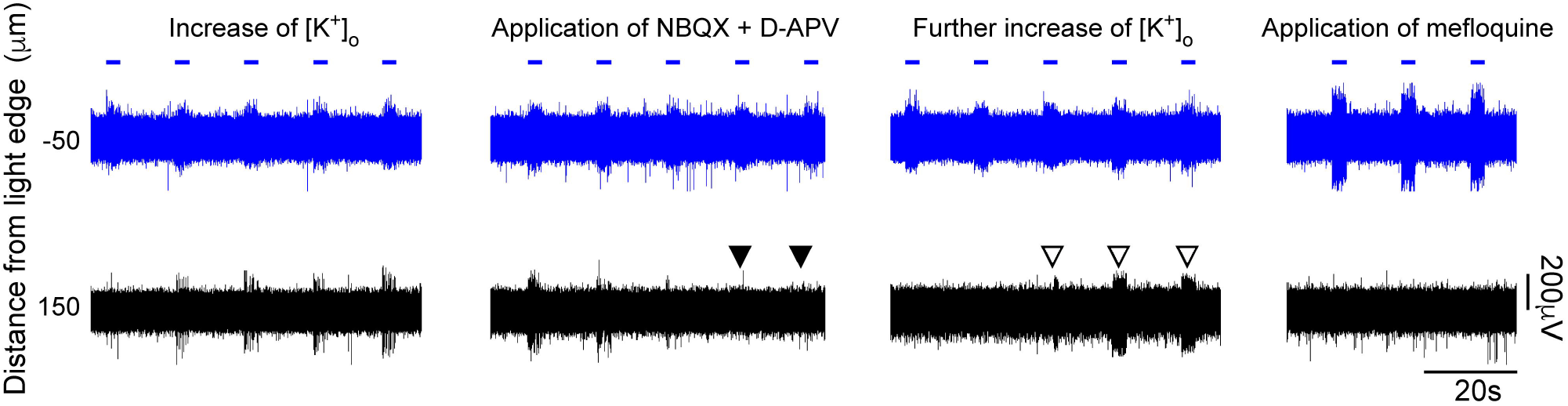
Propagating activity is facilitated by, but is not dependent on, glutamatergic transmission. Repeated propagation assays in the same slice under different pharmacological conditions. In this example, we first saw propagation at 4.5mM [K^+^]_o_ but this was suppressed by the addition of glutamatergic blockers (filled arrowheads). Propagating events then resumed once [K^+^]_0_ was increased further to 5.5mM (open arrowheads). The propagating activity was subsequently blocked entirely by mefloquine.

We next assessed the effects of gap-junction blockers. Unfortunately, this class of drug shows off-target effects (Rozental *et al*., 2001; Srinivas *et al*., 2001; Cruikshank *et al*., 2004), so we examined three different gap-junction blockers: mefloquine, quinine and carbenoxolone. In 4 out of 14 recordings, propagation persisted 15 minutes after the application of a gap junction blocker. In the remaining recordings (10/14 recordings = 71.4%), propagating activity was abolished (see typical examples in Figs. 4–6), and all three drugs showed this effect (quinine, n = 3; mefloquine, n = 6; carbenoxolone, n = 1). The example in Fig. 4 shows the effect of quinine, but this pattern appears representative of the other drugs too, illustrating a rapid and marked decrease in both the amplitude and the firing rate of the recorded activity at electrodes away from the illumination site. Importantly, quinine (unlike the other drugs) can be washed out of the bath (Srinivas *et al*., 2001), and when this was done, propagating activity was rapidly re-established (Fig. 6). When a different gap-junction blocker, mefloquine (see Fig. 5 and 6, right panel), was then applied, this drug also blocked activity propagation. This recording further illustrated another principle, that glutamatergic activity did not sustain this pattern of propagation in the absence of gap-junction coupling.

**Figure 6.**
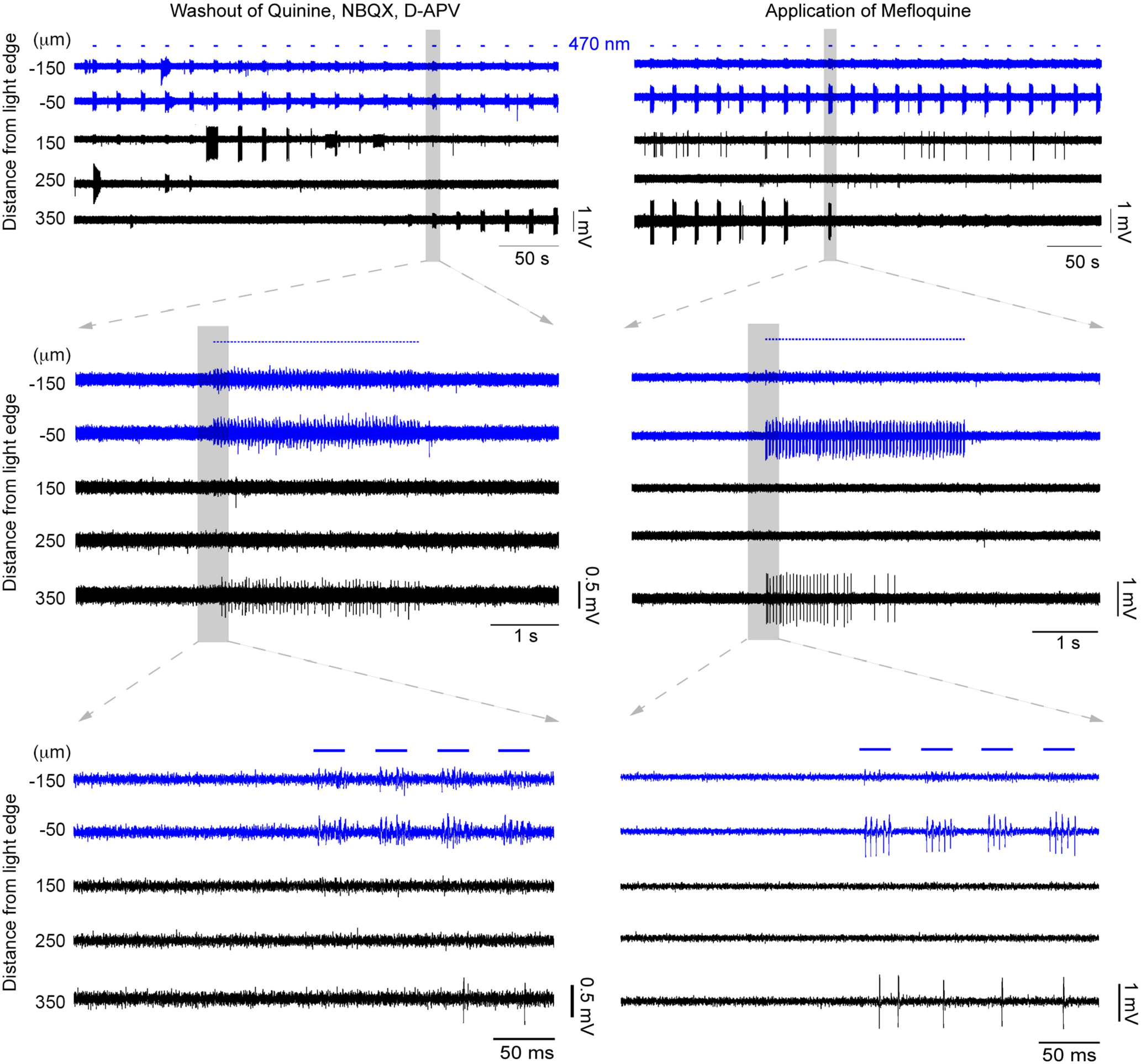
Propagating activity is prevented by multiple different gap-junction blockers. The propagating activity silenced in Fig. 4 was recovered by washing out the blockers (left panel). Stable propagating activity with increasing amplitude was recovered at the electrode 350 μm away. This activity remained strong for several minutes until another gap junction blocker was applied, namely mefloquine (right panel). The propagating activity was once again blocked validating the involvement of gap junctions by using two different blockers.

### Propagation involves primarily fast-spiking cells

Gap junctions in PV-expressing interneurons are believed always to connect only to other PV interneurons (Galarreta & Hestrin, 1999; Hestrin & Galarreta, 2005). We reasoned therefore that analysis of unit spike shapes in the distant electrodes away from the site of stimulation would provide another test of how these events propagate: events that propagate only through gap-junction coupling would show only PV firing at a distance, whereas those that propagate by synaptic means, including glutamatergic or excitatory GABAergic effects would involve a large degree of pyramidal activation. Spiking in PV interneurons has a highly characteristic signature in the extracellular field potential, with narrow spike widths and prominent overshoot, allowing them to be readily distinguished from so-called “regular-spiking” pyramidal cells.

We separated the 20 brain slices that showed extended propagation into three groups: those that were blocked by glutamate blockers (n=6), those that were blocked by gap-junction blockers (n=10), and those that persisted after these pharmacological manipulations (n=4). We analysed the spike waveform of the propagating activity for all three groups. An amplitude-based spike sorting procedure was applied to filter out any background activity before analysing the spike waveform of the time-locked propagating activity. Two features were extracted from each spike waveform: its spike width from valley to peak (measured in ms) and the amplitude ratio between valley and peak (see lower panels in Fig. 6A (Peyrache *et al*., 2012)). These features are typically used to distinguish activity between fast-spiking and regular-spiking cells (Peyrache *et al*., 2012). Our spike waveform revealed 4 putative regular-spiking cells and 16 that had a fast-spiking waveform across all groups (Figure 6A).

Notably, in all the brain slices that were blocked by the gap-junction blockers (group 2), the units were invariably fast-spiking. Pyramidal cells are far more populous than PV interneurons, but on the other hand, usually fire at lower rates. If one assumes that these two effects cancel out, and that therefore one has an equal probability of “finding” a fast-spiking interneuron and a regular spiking neuron, then the probability of finding just fast-spiking interneurons in every single case (n = 10) is approximately 0.1%. In spike sorted data from human neocortex, the ratio of regular to fast-spiking cells is about 80:20 (Peyrache et al., 2012)), in which case, the probability of our result is orders of magnitude lower. We concluded therefore that our finding only fast-spiking neurons, in every brain slice, would only have happened if the propagation were restricted to that cell class, consistent with propagation through the cell-class specific network created by gap-junction coupling.

We then analysed the activity recorded across the multielectrode array with respect to the pulsed timing of the photostimulation (peri-stimulation spike histograms, Fig. 6B), in order to assess the propagation speed of the travelling wave of activity. The speed was calculated from the average latency of the first spikes at distal electrodes. We restricted our analyses only to those experiments where we had pharmacological confirmation of the involvement of gap-junctions (i.e. those using quinine, mefloquine or carbenoxolone). These had a median propagation speed of 59.1mm/s and a spatial extent of up to 0.55mm (see Fig. 6B inset).

### Simulating propagations through the PV-syncytium

The results presented above demonstrate that propagation through the PV-syncytium under conditions of raised extracellular K^+^ is qualitatively different from that induced by 4-aminopyridine (Louvel *et al*., 2001; Gigout *et al*., 2006a; Gigout *et al*., 2006b). In raised [K^+^]_o_, propagation was significantly faster (59.1 mm/s) than in 4-AP (15mm/s) (Gigout *et al*., 2006a). There were further differences in the characteristics of the local field potential: in 4-AP, activity had a large low frequency component, indicative of synchronous activity of many cells, and a broad wave-front (Louvel *et al*., 2001; Gigout *et al*., 2006a; Gigout *et al*., 2006b). In high [K^+^]_o_, on the other hand, propagation is manifested as unit activity (single action potentials from isolated cells). As such, the spread of activity in raised [K^+^]_o_ is more sparse, and is prone to failures of propagation, as evidence showed by the reduced extent of the propagating activity in this model (up to 0.55 mm), compared to 4-AP (> 2mm (Gigout *et al*., 2006a)).

In order to understand these differences better, we developed simulations of biophysically detailed cells that are connected through gap junctions in a 3-dimensional virtual slice. In particular, we were keen to assess how the difference in the cellular electrotonic properties in the two cases impacted on the spread. In both cases, neurons are depolarised relative to baseline, but for different reasons: in high [K^+^]_o_, because the K^+^ reversal potential is relatively depolarised, and in 4-AP, because there is a reduced K^+^ conductance. A further difference is that in 4-AP, the blockade of K^+^ channels will additionally reduce the membrane conductivity, meaning that neurons are more electrotonically compact, thus further facilitating spread through the gap-junctions.

We modelled each neuron as a simple, three compartment model, with a soma and two 200μm-long dendrites (see Fig. 7A). Seventy of these cells were uniformly scattered in a 3-dimensional virtual slice, and randomly connected through gap junctions located on their dendrites (Fig. 7A-B). Each cell had a ‘left’ and ‘right’ dendrite, and its connectivity with the rest of the network was dictated by its location on the x-axis, such that each dendrite only connected to other neurons on that side. This imposed directionality to the network, so when the left part of the virtual slice is stimulated, the activity propagated from left to right. The cell index increased linearly from the side of stimulation (left, id = 1; right, id = 70). The connectivity matrix in Fig. 7C shows an example of a randomly connected network, that follows the connectivity rule in Fig. 7B, and shows that the leftmost cells do not have a direct connection with the rightmost cells.

**Figure 7.**
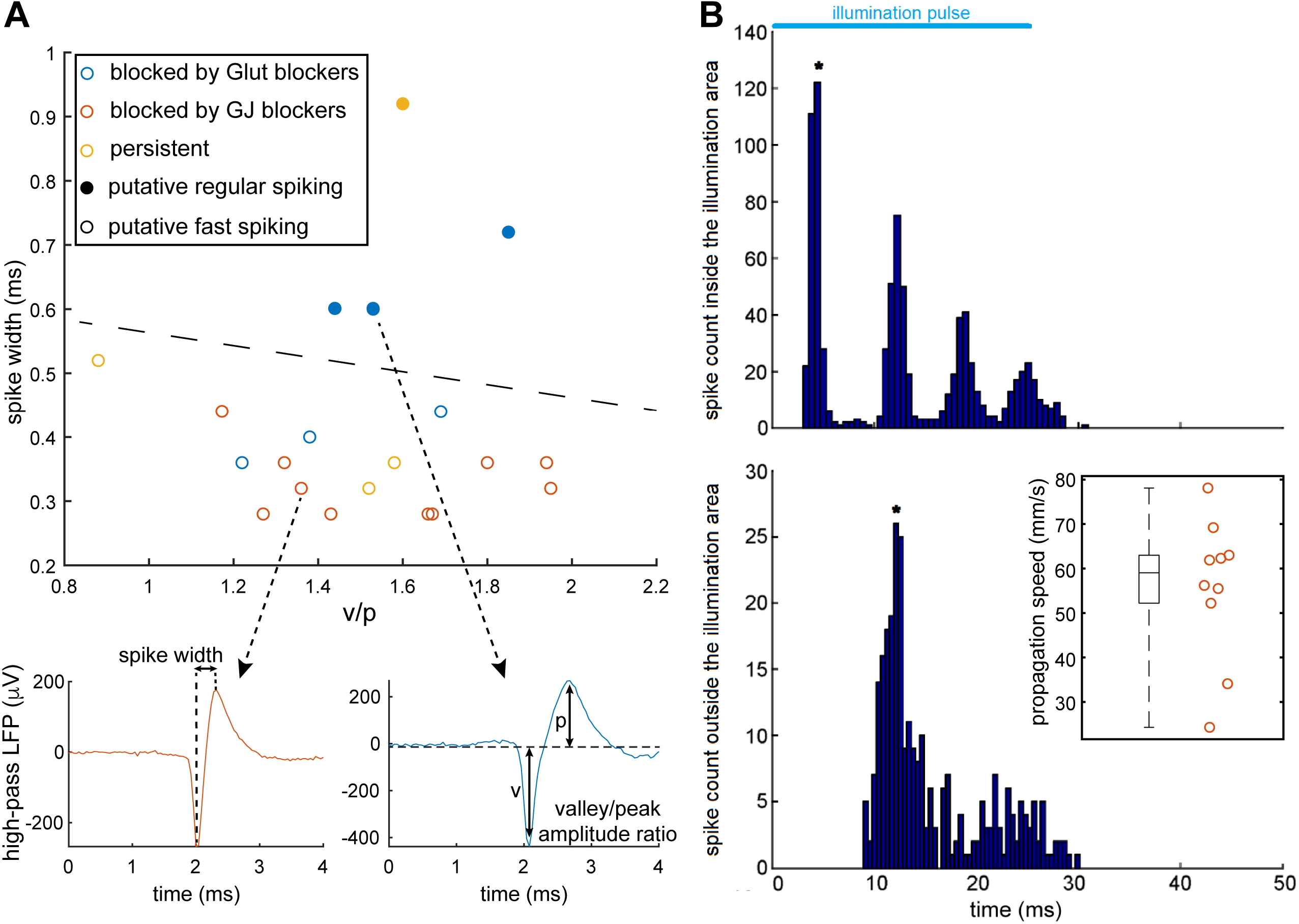
Spike waveform analysis and calculation of propagation speed. (A) The spike waveforms of all 20 pharmacologically manipulated propagations were analysed in terms of their spike width and their amplitude ratio between valley and peak. Both regular and fast spiking waveforms were observed (see lower panels for average spike examples). Notice that none of the hypothesized gap-junction mediated activity was found to exhibit a regular-spiking waveform (i.e., spike width > 0.55ms). (B) Example analysis of the speed of propagation calculation, for the gap-junction mediated propagations. The spike time histograms from two different electrodes were plotted: one inside the illumination area (close to the border) and one outside the illumination area where propagating activity was detected. The period covered by the histogram matches the period of the illumination (50ms). The first peak in each histogram is marked by asterisk. The propagation speed was directly calculated from the time difference between these peaks and the distance between the respective electrodes. The inset shows the distribution of the calculated propagation speeds (only for propagations blocked by gap junction blockers). It reveals high variability (standard deviation 17.2 mm/s) and a median speed of 59.1 mm/s.

Cells located in the left-most 200μm of the virtual slice were stimulated 50ms after starting the simulation, and the stimulation pulse, delivered directly to their soma, lasted for 25ms. The exact number of cells stimulated varied slightly from simulation to simulation since the scattering of the cells in the virtual slice was random. Simulations were run for each of the three cases: (1) the control case, where the settings are set to default; (2) the high K^+^ case, where the extracellular potassium concentration is considered to be raised to 10.5mM, instead of the default 3.5mM, thus changing E_K_ and also therefore, raising the resting membrane potential, and reducing the effective action potential threshold; and (3) the 4-AP case, where the membrane resistance is five times higher than the default value and the K^+^ channels are almost entirely blocked (conductance is set to 2% of their default value). Typical results of these simulations are shown in Fig. 7D for each one of the different cases. The blue regions in the scatter plots represent the temporal and spatial extend of the stimulation. For the control case, only the cells in the stimulation area were active, and this activity did not spread beyond the stimulation area at all. In the high [K^+^]_o_ case, activity propagated beyond the area of stimulation, but typically failed before the end of the slice (i.e., 70^th^ cell). Furthermore, we found that even within the propagating territory, the wave of activity could skip some neurons, since there are multiple paths across the network. Thus, not every cell participates in the propagation. Interestingly, the propagation did not advance smoothly. Rather, following stimulation, there was a rapid and almost simultaneous activation of a group of cells in the middle of the slice, followed by a delay before the next set of neurons were recruited.

Gap-junction mediated propagation in the 4-AP simulations, though, were qualitatively different. There was a more gradual and slow propagation that reliably reached the end of the slice without failing. The participation of the cells was complete, with few, if any, being skipped during the propagation (median = 100% as opposed to 71.1% for high [K^+^] simulations; p << 0.001, two-sided Wilcoxon rank sum test; Fig. 7F). The distribution of the propagation speeds in 200 simulations for each case is shown in Fig. 8E. The median propagation speed for the high K^+^ case is 57.1 mm/s. This value is close to what was expected from the experimental results presented above. The median propagation speed of the 4-AP case is significantly lower (33.5 mm/s; p << 0.001, two-sided Wilcoxon rank sum test) but higher than the expected one from previous studies (15mm/s) (Gigout et al., 2006a).

**Figure 8:**
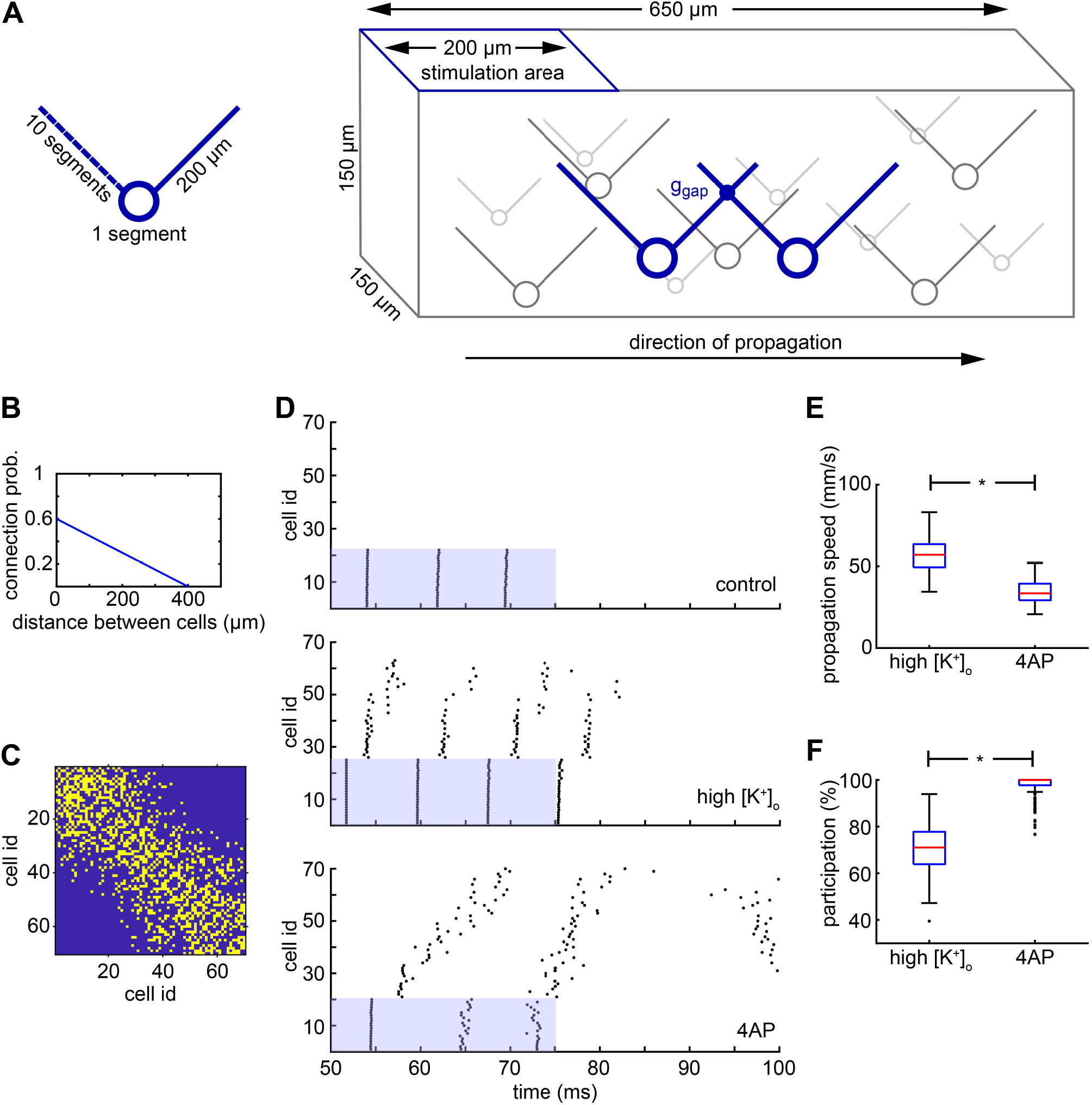
Simulation of the high K+ propagation and comparison with the 4-AP propagation. (A) Schematic of the model cell and of the model network of electrically connected cells. The cell has a simple morphology with a soma and two dendrites, one on the left and one on the right. The cells are randomly placed in a 3-dimensional virtual slice and they form a network through gap junctions. The left dendrite for each cell is used to connect it with other cells located on its left whereas the right dendrite is used to connect it with cells on the right. The left side of the network is stimulated and the activity propagates to the right. (B) The probability of connection between two cells is linearly decreased as the distance between them increases. (C) Example of a connectivity matrix of a randomly generated network where the yellow colour indicates a connection. The cell id is derived from the ordering of the cells on the x-axis, from left to right. Notice that the leftmost cells do to have any direct connection with the rightmost cells due to the limited length of their dendrite (200 μm). (D) Typical results of the simulation under three conditions: control, high K+, and 4-AP. In the control case there is no propagation; only the cells in the stimulation area fire. In the high K+ case, there is a fast propagation to the right side of the network but not all cells are participating. In the 4-AP case, there is qualitatively different propagation where the speed is lower but almost every cell in the network participates. (E) Distribution of propagation speeds for the high K+ and 4-AP cases. The propagation under high K+ conditions is significantly faster (p << 0.001, two-sided Wilcoxon rank sum test). (F) Distribution of participation percentages for the high K+ and 4-AP cases. The participation under 4-AP conditions is significantly higher (p << 0.001, two-sided Wilcoxon rank sum test).

## Discussion

In these studies, we demonstrate propagating waves of activity within the population of fast-spiking interneurons, extending at distance from the point of onset, at levels of [K^+^]_o_ that are commonly seen during a seizure (Somjen, 2004). Previous work has only shown gap-junction coordination very locally, through directly connected cells (Galarreta & Hestrin, 1999; Gibson *et al*., 1999). Notably, we were able to identify a threshold level of [K^+^]_o_ around 8mM for this propagating activity pattern; below that, at physiological levels of [K^+^]_o_, activation of PV interneurons remains very focal. Previously, several groups have demonstrated the coordination of interneuronal activation across single gap-junctions (Galarreta & Hestrin, 1999; Gibson *et al*., 1999), but for more extensive propagation within cortical networks, gap junction-facilitated propagation over extended distances has only been demonstrated using pharmacological manipulation, bathing tissue in 4-AP (Szente *et al*., 2002; Gajda *et al*., 2003; Gigout *et al*., 2006a). This pharmacological manipulation, while of interest, does represent a rather extreme disruption of neocortical interneuron behaviour (Codadu *et al*., 2019a). Of particular relevance to the present study is that 4-AP makes neurons more electrotonically compact, by blocking a large component of K^+^ conductance, and will thus naturally facilitate electrotonic propagation. Our new data is the first to show spatially extended propagation in conditions that are known to occur naturally in the living brain.

Since 4-AP has a disproportionately large effect on the population of parvalbumin-expressing interneurons (Codadu *et al*., 2019a), we also chose to study the effect of raised [K^+^] in this same population. Electrical coupling between this population of interneurons is well established (Galarreta & Hestrin, 1999; Gibson *et al*., 1999), but coupling has also been described for other populations of cortical interneurons (Galarreta & Hestrin, 2001; Hestrin & Galarreta, 2005), and our findings are likely to generalise to these interneuronal populations too.

One notable finding is that the pattern of gap-junction coupled propagation in high [K^+^]_o_, where we see tightly time-locked and rapid propagation of single action potentials, appears different from that in 4-AP, which takes the form of a broad wave-front, with a relatively small high frequency component (e.g. (Gigout *et al*., 2006a; Gigout *et al*., 2006b; Parrish *et al*., 2018). A further distinction between these two experimental paradigms is that 4-AP waves additionally involve other interneuronal populations, and may also trigger waves of raised [K^+^] themselves, secondary to GABAergic induced chloride-loading (Viitanen *et al*., 2010), both of which would be expected to broaden the wave. It is noteworthy therefore that 4-AP also blocks K^+^ channels in glia (Hosli *et al*., 1981), which may boost the transient rise in [K^+^]_o_, an effect which may be further enhanced by gap-junction blockers. The resultant slow transient of [K^+^]_o_ following a protracted burst of interneuronal activity (Viitanen *et al*., 2010) may also contribute to the broad wave-front in 4-AP.

Extended propagation of activity within the PV population only occurred at raised levels of [K^+^]_o_, but appears also to be boosted by background levels of excitatory neurotransmission. Blockade of glutamatergic neurotransmission led to a failure of propagation in a proportion of brain slices, but interestingly, it was possible to resurrect the propagating events by further raising the bathing [K^+^]_o_. Together these data support the model we present, in which the likelihood of propagation is dictated by level of depolarisation of the cell at rest, which dictates the ease with which the next element in a chain of neurons is recruited, and how electrotonically compact the network is. Both raised [K^+^]_o_ and a tonic level of glutamatergic drive facilitates propagation by the first mechanism, while blocking K^+^ conductances using 4-AP facilitates it by making each element more electrotonically compact. It is possible that excitatory GABAergic activity could also facilitate propagation in this same way, although we did not test this explicitly. It is important though that our results are not consistent with propagation solely through excitatory GABAergic effects, because this would lead to non-specific activation of different cell classes, whereas our spike-sorting analyses indicate that in the great majority of cases (and all cases in which propagation was suppressed by gap junction blockers), propagating activity is restricted to the fast-spiking class of interneurons. Thus, any contribution from excitatory GABAergic effects is likely only to be an adjunct to the coupling we observed, rather than the primary means of propagation.

The primary effect of enhancing the electrical coupling of interneurons, in this way, will be to extend the inhibitory surround during focal cortical activation (Prince & Wilder, 1967; Schwartz & Bonhoeffer, 2001; Trevelyan *et al*., 2006; Parrish *et al*., 2019), while also enhancing coordination of this inhibitory effect. Interneuronal activity at this time, however, is a double edged sword, because there is now good evidence that these neurons may become active drivers of epileptic discharges, if pyramidal cells become loaded with chloride (Huberfeld *et al*., 2007; Dzhala *et al*., 2010; Avoli & de Curtis, 2011; Ellender *et al*., 2014; Pallud *et al*., 2014; Alfonsa *et al*., 2015; Raimondo *et al*., 2015; Alfonsa *et al*., 2016; Chang *et al*., 2018; Magloire *et al*., 2019). Raised extracellular K^+^ will also facilitate chloride loading, which is coupled via the cation-chloride co-transporter, KCC2. In either case, whether GABAergic output is inhibitory or excitatory, the enhanced coupling of interneurons through their gap-junctions is likely to be an important determinant of the complex pattern of propagating local field potentials during a seizure (Viventi *et al*., 2011; Smith *et al*., 2016; Codadu *et al*., 2019b). It is also highly pertinent that gap-junction expression is commonly increased in epileptic brains (Manjarrez-Marmolejo & Franco-Perez, 2016). Whether this is protective, or serves to exacerbate the epileptic condition, remains an open question.

## Abbreviations

PV: Parvalbumin
ACSF: artificial cerebro-spinal fluid
NMDA: N-methyl-D-aspartate
AMPA: α-amino-3-hydroxy-5-methyl-4-isoxazolepropionic acid
APV: amino-5-phosphonovaleric acid
4-AP: 4-aminopyridine

